# UALCAN Mobile, an app for cancer proteogenomic data analysis

**DOI:** 10.1101/2025.09.04.674198

**Authors:** David Rubey, Darshan Shimoga Chandrashekar, Ahmedur Rahman Shovon, Gopi Chand Puli, Santhosh Kumar Karthikeyan, Upender Manne, Chad J. Creighton, Sidharth Kumar, Sooryanarayana Varambally

## Abstract

Cancer is a complex disease affecting various organs and is a major cause of death worldwide. During cancer initiation, disease progression, and tumor metastasis, various genomic and proteomic alterations are observed. Recent technological advances have led to the generation of large amounts of molecular data, including genomics and transcriptomics. These large-scale datasets can be utilized to analyze and identify sub-class-specific cancer biomarkers and targets. However, there is a need for the development of user-friendly tools for large-scale data analysis, disseminating the analyzed data in a visualizable format to cancer researchers with no programming skills. We developed UALCAN, a comprehensive platform that allows users to integrate disparate data to better understand the genes, proteins, and pathways perturbed in cancer and make discoveries of potential biomarkers and targets. In the current study, we describe the development of the UALCAN Mobile application (app) that will provide cancer transcriptomic data obtained from The Cancer Genome Atlas (TCGA) project to evaluate protein-coding gene expression based on various stratifications, including stage, grade, race, gender, and molecular-subtypes across over 30 types of cancers. In addition, the UALCAN mobile provides data analysis options for epigenetic changes due to DNA promoter methylation and Clinical Proteomic Tumor Analysis Consortium (CPTAC) cancer proteomic data. The app provides access to large cancer molecular datasets on the go. To find changes in the expression of causative genes and proteins and to identify biomarkers and therapeutic targets, UALCAN mobile app will be extremely valuable. The “UALCAN Mobile” app is free to use and can be downloaded from both the iOS/Apple and the Android Play Store and has been downloaded over 100 times in each of iOS and android app stores.

**Key Points:**

- Cancer, a complex disease affecting various organs, is a major cause of death worldwide.
- Many molecular alterations, gene and protein expression changes lead to the initiation of cancer, disease progression, and tumor metastasis.
- The available large genomic and proteomics datasets can be utilized to analyze and identify sub-class-specific cancer biomarkers and targets with innovative tools.
- UALCAN Mobile application (app) will provide gene expression, promoter DNA methylation, and protein expression across cancers based on various stratifications, including stage, grade, and molecular subtypes across over 30 types of cancers.
- UALCAN Mobile app is free and can be downloaded from both iOS/Apple and the Android Play Store.

## Introduction

Cancer, a complex and heterogeneous disease, shows various molecular alterations in genes and proteins that occur during disease progression. Identifying new biomarkers and therapeutic targets is essential for enhancing precision targeting. Recent developments in high-throughput technologies enabled large-scale genomic, transcriptomic, and proteogenomic data generation across cancer types. The availability of these data and the ability to analyze them using bioinformatics have enabled the discovery of many genomic aberrations and differential gene expression patterns in various cancers and during disease progression. The provision of readily accessible analysis platforms has allowed clinicians and cancer researchers to discover and validate cancer biomarkers. To facilitate basic to detailed analysis of the large cancer datasets, the research community has developed various web-based/standalone tools. One popular tool, cBioPortal (1, 2), allows users to submit sets of genes for a cancer type of interest and query for genomic and transcriptomic aberrations. cBioPortal mainly focuses on gene mutation and copy number alterations (CNA) data and provides visualization highlighting mutation and CNA patterns and gene expression for specific genes. Other analysis and visualization portals such as miRGator v3.0 (3), EpiCompare (4), TANRIC (5), ISOexpresso (6), Zodiac (7), MiPanda (8), and integrated genome browser (IGB) (9,10) provide analysis and visualization options for specific biomolecules. Our earlier efforts resulted in the development of a platform for comprehensive cancer data analysis, UALCAN, a web-based tool that enables researchers to access omics data and to perform integrative data analyses (11,12). UALCAN is a comprehensive and user-friendly web portal for analyzing cancer transcriptome data. It provides various tools and visualizations to help researchers explore gene expression patterns across cancer types and corresponding normal tissues.

As the cancer research landscape evolves, there is a growing demand for more portable and convenient access to these valuable resources. Although in 2017, a mobile application called GE-mini was released, it lacked an in-depth tumor subtype analysis feature (13). Thus, we realized an opportunity to further disseminate the data to mobile devices, including phones and tablets. With this goal, we developed UALCAN Mobile, a freely downloadable app for cancer transcriptome data analysis using TCGA data. It distinguishes itself from the UALCAN website primarily through its mobile-centric design and enhanced user experience tailored for smartphone and tablet usage. Designed as a hybrid mobile application, it can be downloaded and installed on smartphones and tablets (Android and iOS), providing convenient access to cancer genomics data even in mobile environments. UALCAN Mobile allows researchers to access RNA-seq data from The Cancer Genome Atlas (TCGA) and to perform gene expression analysis of protein-coding genes across over 30 types of cancers. Furthermore, UALCAN mobile provides DNA promoter methylation data for analysis, which is known to regulate protein-coding gene expression. To provide changes in protein expression in cancer, UALCAN mobile provides stratified data from the Clinical Proteomic Tumor Analysis Consortium (CPTAC). UALCAN Mobile also allows analysis of subgroup-specific gene expression. The data, figures, and statistics of the analyses are downloadable. To enhance the utility of this app, it will, in the future, include patient survival and ChIP-sequencing data.

## Methods

### Gene expression, promoter methylation, and proteomic data

The TCGA data was collected using TCGA-Assembler (14), and used to download TCGA level 3 RNA-seq data related to over 30 cancer types. It was installed on R 3.2.2 (https://cran.r-project.org/). Using TCGA assembler, a) “rsem.genes.results” files were obtained for ‘Primary Solid Tumor’ and ‘Solid Tissue Normal’ for each cancer type and b) DNA methylation data generated via Illumina HumanMethylation450 array for all tumor and normal samples were collected. In addition, we downloaded protein expression data generated using high-throughput mass spectrometry by the Clinical Proteomic Tumor Analysis Consortium (CPTAC) for 17 cancer types. Detailed methodology of data collection and analysis has been provided in the articles describing the UALCAN web platform (11,12). The downloaded gene/protein expression data and DNA methylation profiles were further stratified based on various parameters such as age, sex, race, survival status, tumor grade, and tumor stage, among others, for each patient in the XML (eXtensible Markup Language) file. A PERL script was written to parse all XML files corresponding to a specific cancer and extract them into a tab-separated file.

### Development of the UALCAN Mobile App

The UALCAN Mobile app is built upon a foundation of modern web technologies, allowing the use of its functionalities across various platforms. The core framework for the app’s user interface is Angular (https://v15.angular.io/docs, v15.0.0), a popular JavaScript framework known for its single-page application architecture. This is further enhanced by the Ionic framework (https://ionicframework.com, v6.3.9), a toolkit designed to build hybrid mobile applications. The combination of Angular and Ionic allows for a single codebase to be utilized for the app’s functionalities, streamlining development and ensuring consistency across platforms. TypeScript (https://www.npmjs.com/package/typescript/v/4.8.2, v4.8.2), a superset of JavaScript, provides the app with the benefits of static typing and improved code maintainability. The app leverages the capabilities of the Highcharts library (https://www.highcharts.com) to generate box plot charts of gene expression. These charts effectively communicate the distribution of gene expression levels across various sample groups. A key aspect of the UALCAN Mobile app is its capability to function as a hybrid mobile application. Apache Cordova (https://cordova.apache.org) acts as a bridge, facilitating the compilation of the app’s single codebase into platform-specific versions for iOS and Android devices. This approach allows users to access the app’s functionalities regardless of their preferred mobile operating system. Several additional dependencies contribute to the UALCAN Mobile app’s functionality. The Angular Material library (https://material.angular.io, v15.0.3) provides pre-built UI components that adhere to Google’s Material Design guidelines, enhancing the app’s visual appeal and user experience. Ionic Native plugins, such as those for core functionalities (ionic-native/core, v5.36.0), file access (ionic-native/file, v5.36.0), and PDF generation (ionic-native/pdf-generator, v5.36.0), bridge the gap between web-based development and native device functionalities. The UALCAN Mobile app retrieves data from the UALCAN application-programming interface (API), which acts as the backend for the application. The API prepares data using a PERL CGI script hosted on UALCAN’s web server, maintained by the UAB Department of Pathology. This API securely transmits data in JSON format to the mobile app over HTTPS, enabling the visualization of box plot for gene expression.

### Design decisions

The UALCAN Mobile app’s architecture and implementation are guided by a series of design decisions aimed at ensuring a robust, user-friendly, and cross-platform experience for cancer data exploration. App architecture to support cross-platform compatibility: The UALCAN Mobile app’s architecture is designed to achieve seamless cross-platform compatibility. The design decision of using Angular with the Ionic stack enables a single codebase targeting both IOS and Android platforms. Compilation with Apache Cordova provides near native performance and unified updates across platforms. The use of TypeScript ensures type safety and code reusability.

All cancer omics data is served through a secure RESTful API using PERL CGI, with responses in JSON over HTTPS. Each query retrieves all relevant stratifications in a single API request with the support of client-side filtering to reduce network overhead. This design minimizes latency, lowers API load, and enables offline analysis of previously fetched data. As mobile devices have limited computational power, the fetched data is visualized using mobile-friendly Highcharts. The app renders box plots and statistical summaries directly from API payloads, supporting user-driven exploration with minimal UI lag. To support the native device features such as file storage and PDF export, UALCAN Mobile uses Ionic Native plugins. A core challenge in visualizing gene expression data on mobile devices is ensuring clarity and usability across diverse screen sizes and both landscape and portrait orientations. To address this, we implemented responsive layouts and dynamic chart resizing within the app, allowing complex visualizations to render effectively regardless of device or orientation. Highcharts’ adaptive features, along with Google’s material design, provide a smooth user experience and clear visualization for large genomic and proteomics datasets.

## Results

### UALCAN Mobile app architecture

UALCAN Mobile utilizes a single codebase to generate Android and iOS applications as provided in **Figure 1**. Powered by the Ionic framework, the UALCAN Mobile app capitalizes on its versatility and popularity in hybrid app development. By leveraging Apache Cordova, the codebase is compiled into platform-specific versions, ensuring compatibility and performance across diverse mobile devices. Data retrieval in the UALCAN Mobile app is orchestrated securely through the UALCAN API, a PERL CGI script hosted on the UAB server. Data transmission occurs over HTTPS in JSON format, safeguarding the integrity and confidentiality of the information exchanged between the server and the mobile app. This cross-platform approach allows the UALCAN Mobile app to be deployed and accessed on various Android and iOS devices while maintaining a unified codebase.

**Figure 1.**
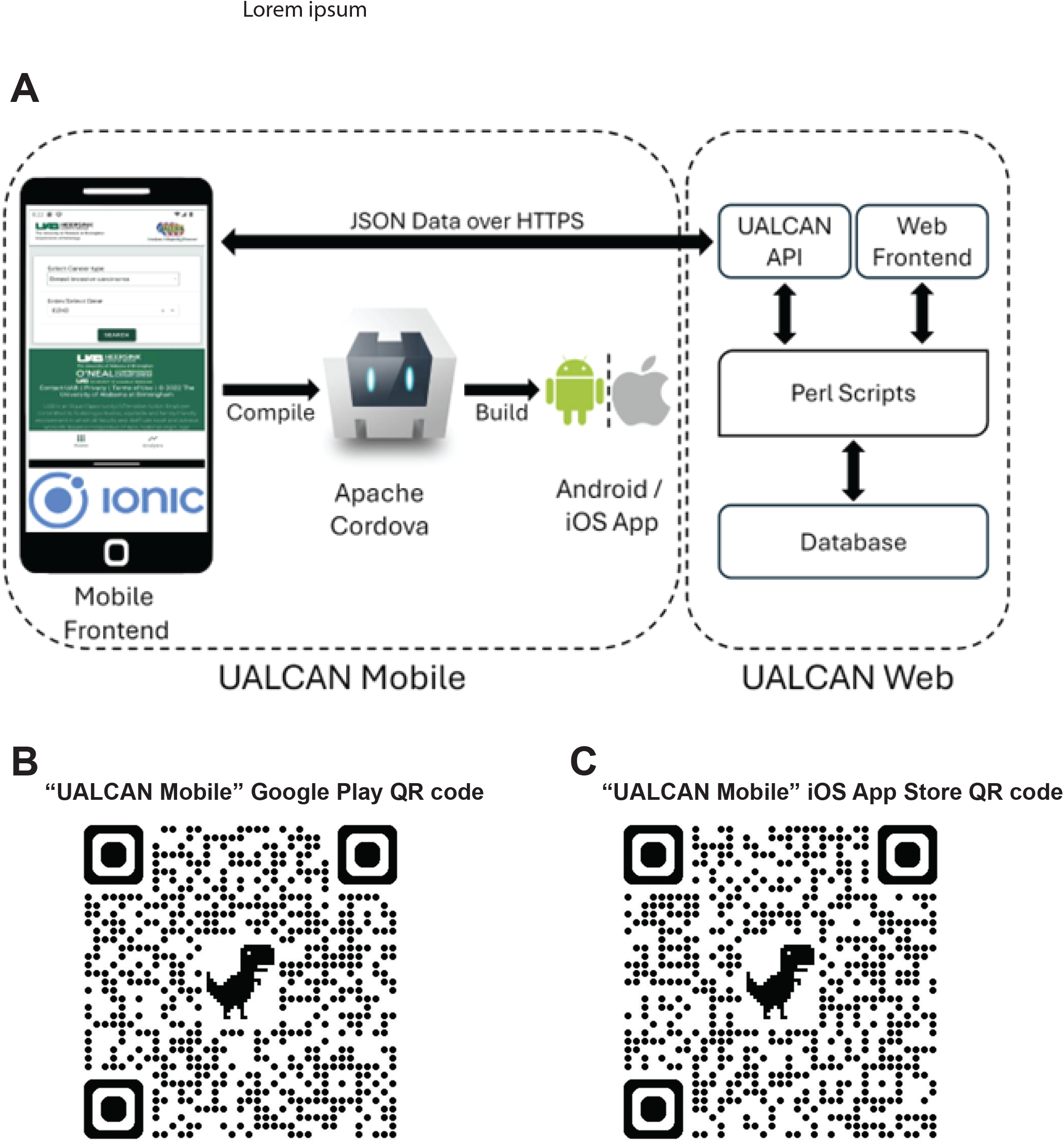
**A. UALCAN Mobile app architecture. B-C**, The QR codes for tablets, Apple, and Android mobile devices.

Balancing efficiency and interactivity in cancer data exploration: The UALCAN Mobile app achieves a balance between minimizing API calls and maintaining a user-friendly, interactive experience. This technical feat is achieved through a data flow architecture that is appropriately designed. Quick Response (QR) codes for tablets, Google Play Store app, and Apple App Store are provided in **Fig. 1B** and **Fig. 1C**.

Key steps and interactions involved in the app’s functionality are provided in **Figure 2**. The user initiates the process by selecting a cancer type and gene of interest within the app. This triggers a single call to the UALCAN API, securely transferring the selections over HTTPS. The UALCAN API then queries its database to retrieve the relevant TCGA RNA-seq data based on the user’s choices. Notably, the API fetches data for all categories (individual stages, gender, and race) at once. This approach minimizes the number of subsequent API calls needed when the user explores the data by filtering through these categories within the app itself. This local processing minimizes network traffic and ensures a smooth, responsive user experience even with limited internet connectivity.

**Figure 2.**
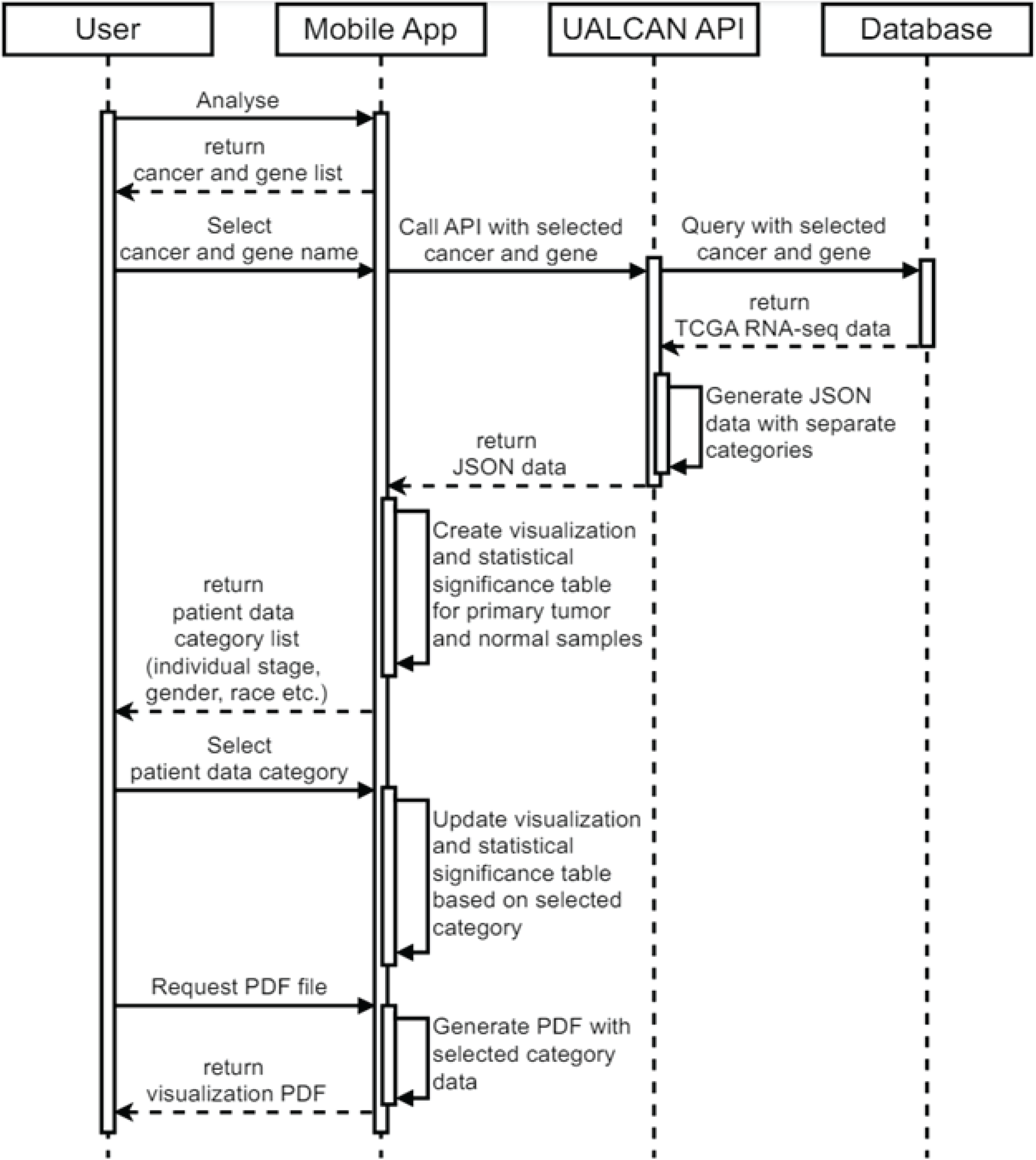
Flow chart of steps and interactions involved in UALCAN Mobile.

### UALCAN mobile app usage

The UALCAN Mobile app can be accessed and downloaded from the Google Play Store (https://play.google.com/store/apps/details?id=edu.uab.path.ualcan) and the Apple App Store (https://apps.apple.com/us/app/ualcan-mobile/id6450547634), fulfilling the requirements outlined. We focused on simplicity in our efforts to create an intuitive application interface for cancer researchers. The home page provides a brief overview of the app. Users can access the analysis section by clicking the hyperlink at the bottom of the page [**Figure 3A**]. The analysis page presents the user with two input fields: a) a drop-down menu to select one of 33 different tumor types and b) a text box to enter the gene of interest [**Figure 3B, Figure 4A**].

**Figure 3.**
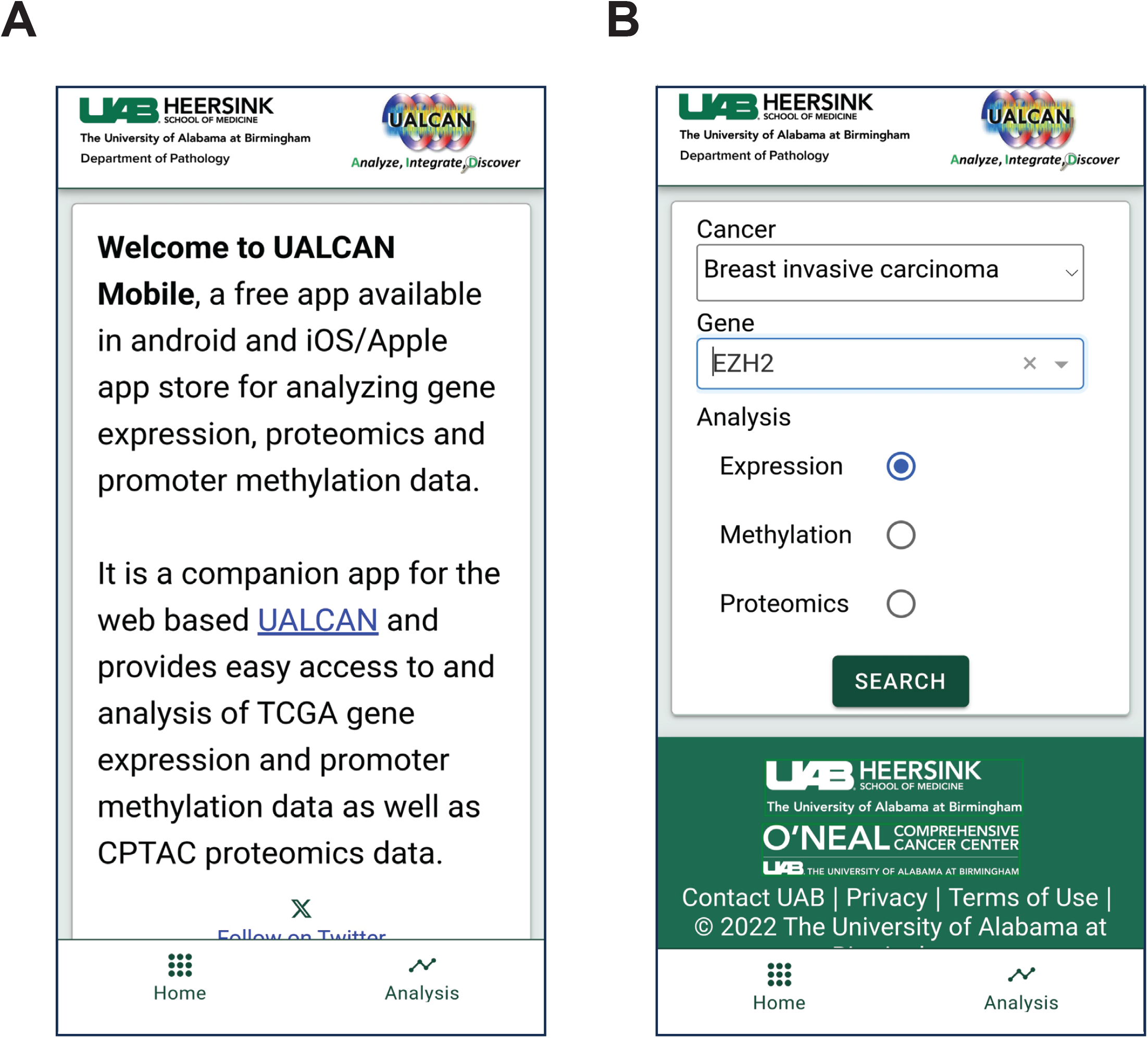
UALCAN mobile app usage. **A**. The app’s opening page. The analysis option is in the bottom right corner. **B**. Selection of cancer type and gene of interest for analysis.

**Figure 4.**
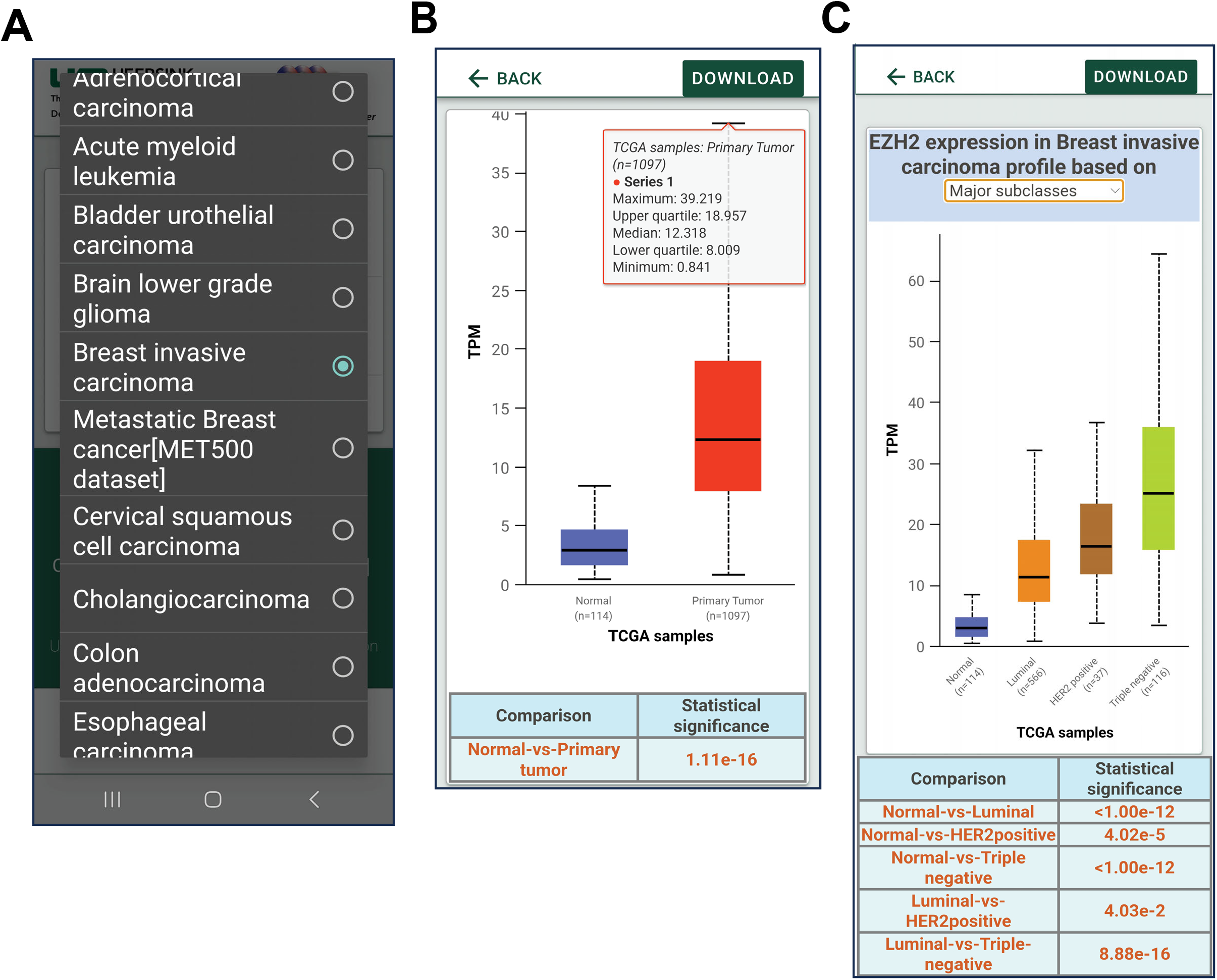
UALCAN mobile app data analysis. **A**. Users can select the type of cancer to be analyzed. **B**. Analysis showing differential expression of the gene *EZH2*. **C**. *EZH2* expression based on breast cancer subtypes.

Upon submitting their query, users gain access to expression patterns for their selected gene. These patterns are available not only for normal and tumor samples but also for various tumor subtypes. Users can refine their analysis based on factors such as tumor stage, patient gender, race, and molecular subtypes [**Figure 4B-C, Figure 5A-B**]. The associated statistics can be visualized at the bottom of each page of analysis. This basic approach to the visualization of large-scale data on cancer gene expression enhances the utility of this app. Similarly, UALCAN Mobile provides information about the changes in the DNA promoter methylation, a type of epigenetic regulation for each of the gene targets and the expression of that gene (**Fig. 6 A-B**). Users can compare the promoter methylation of any genes and associated gene expression using the app for any specific genes across cancers. Researchers can also evaluate the protein expression changes in various cancers using UALCAN Mobile, as it provides incorporated CPTAC proteomics data (**Fig. 6 C-D**). To date, “UALCAN Mobile” app has been downloaded over 100 times in each of iOS and android app stores.

**Figure 5.**
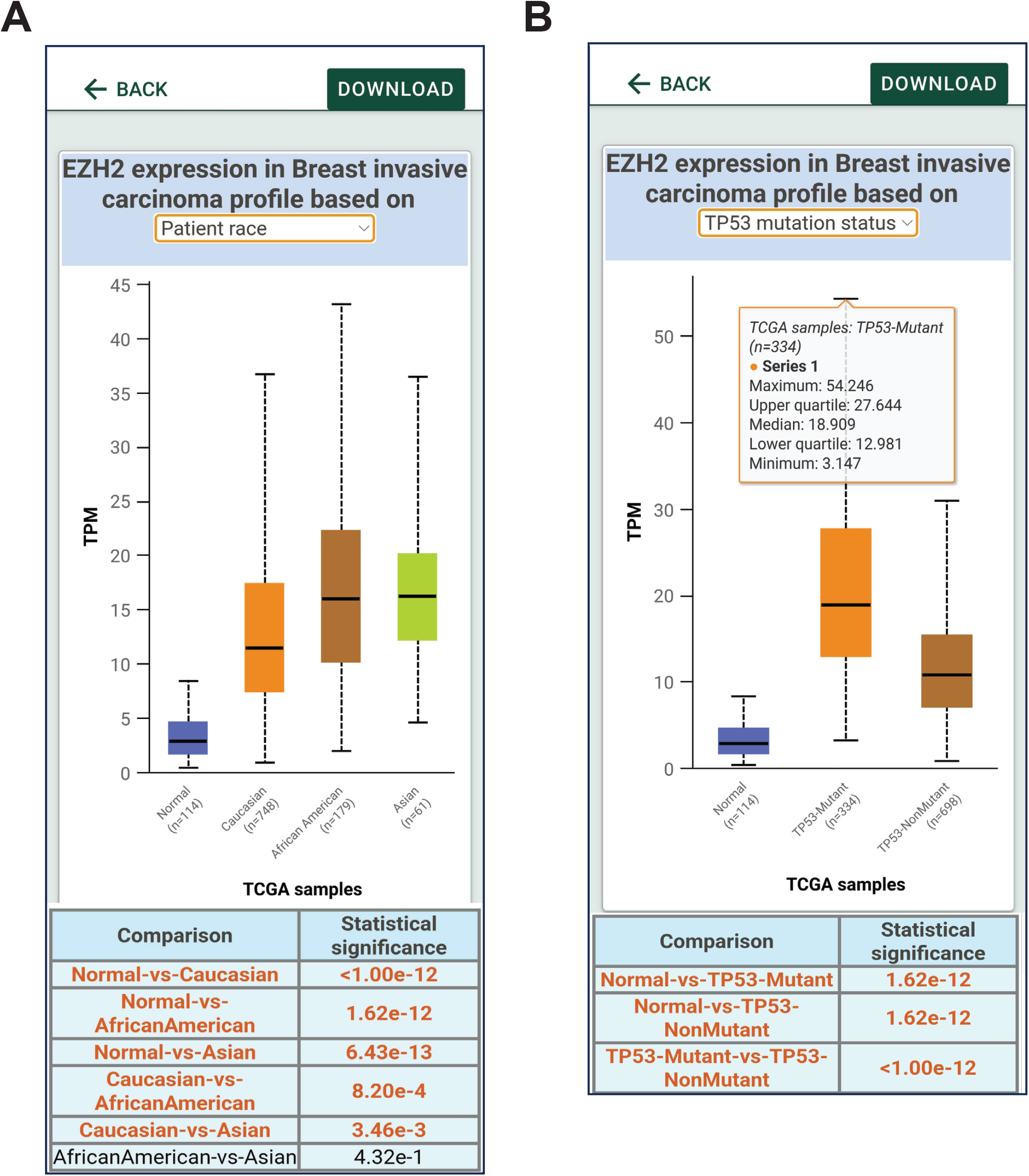
UALCAN mobile app data analysis based on patient stratification. **A**. Gene expression can be stratified based on patient race. **B**. EZH2 expression based on the p53 mutational status.

**Figure 6.**
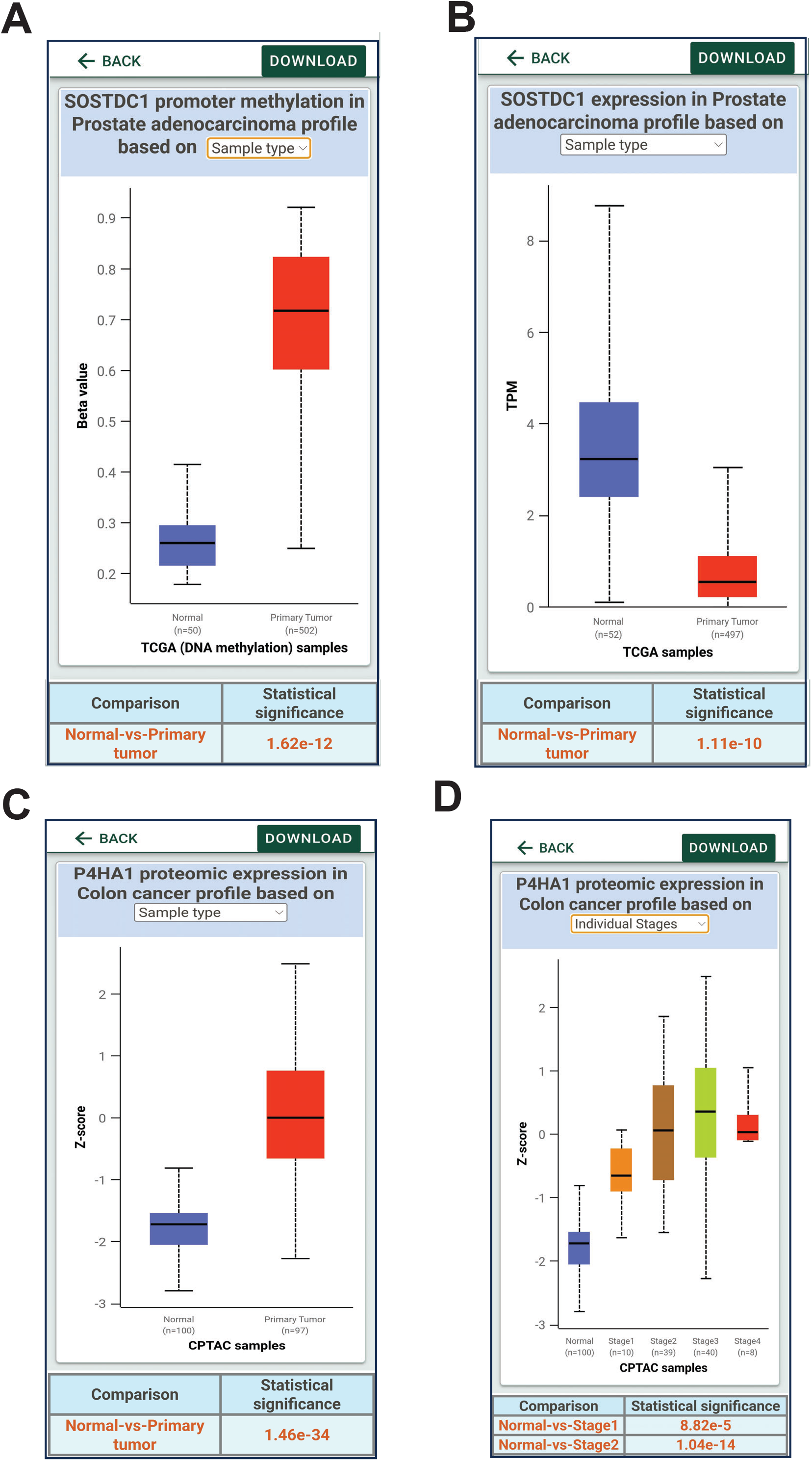
Analysis of promoter methylation and protein expression data using UALCAN mobile app. **A, B**. DNA Promoter methylation of SOSTDC1 gene in prostate cancer data from TCGA and corresponding gene expression. **C**,**D**. prolyl 4-hydroxylase subunit alpha 1 (P4HA1) protein expression in colon cancer patients from CPTAC proteomic data.

## Discussion

As more and more large-scale cancer sequencing and molecular data become available, easy accessibility and analysis are needed to identify cancer biomarkers and targets to tackle the menace of cancer and enhance treatment options. The molecular profiling data from the NCI TCGA consortium is proving to be a great resource for various analyses and patient stratification. Using TCGA data along with other data, including the Clinical Proteomic Tumor Analysis Consortium (CPTAC), we created an integrative cancer data analysis platform, UALCAN. UALCAN enables researchers to identify various biomarkers and perform survival analyses for a gene of interest across multiple cancer types and subtypes. To further enhance and disseminate the data and make it available to the fingertips of users, we embarked on developing “UALCAN Mobile”, a free application that is downloadable on IOS/Apple and Android app stores. Although the app does not provide all the analysis and graphics available on the web platform, it does make the data highly portable once the app is downloaded by the users. We anticipate the use of this cancer data analysis app in many scenarios, including cancer research seminars and conferences, as well as during reviews to confirm or glance at the genes or cancers of interest. In the future, we will enable additional analysis in the app, such as gene co-expression analysis, pan-cancer analysis, and non-coding RNA analysis We anticipate that the UALCAN Mobile app will be a useful companion tool for cancer biologists and clinicians and will aid in the identification of new cancer biomarkers and therapeutic targets. With the large user base that the UALCAN web platform has accrued, we expect the UALCAN Mobile app to become a distinctive app for the analysis of cancer molecular data and provide the data to users worldwide and anytime. The goals are to allow cancer researchers and non-cancer researchers to query any gene of interest and to assess its associations with diseases.

## Acknowledgements

The cancer molecular data used in this app are from the TCGA and CPTAC consortium. This work is supported by the Breast Cancer Research Foundation of Alabama to SV. SV and UM are supported by U54 CA118948. We thank Dr. Don Hill for editing.

## References

1. Cerami E, Gao J, Dogrusoz U, Gross BE, Sumer SO, Aksoy BA, et al. The cBio cancer genomics portal: an open platform for exploring multidimensional cancer genomics data. Cancer Discov 2012;2:401–4

2. Gao J, Aksoy BA, Dogrusoz U, Dresdner G, Gross B, Sumer SO, et al. Integrative analysis of complex cancer genomics and clinical profiles using the cBioPortal. Sci Signal 2013;6:pl1

3. Cho S, Jang I, Jun Y, Yoon S, Ko M, Kwon Y, et al. MiRGator v3.0: a microRNA portal for deep sequencing, expression profiling and mRNA targeting. Nucleic Acids Res 2013;41:D252–7

4. He Y, Wang T. EpiCompare: an online tool to define and explore genomic regions with tissue or cell type-specific epigenomic features. Bioinformatics 2017;33:3268–75

5. Li J, Han L, Roebuck P, Diao L, Liu L, Yuan Y, et al. TANRIC: An Interactive Open Platform to Explore the Function of lncRNAs in Cancer. Cancer research 2015;75:3728–37

6. Yang IS, Son H, Kim S, Kim S. ISOexpresso: a web-based platform for isoform-level expression analysis in human cancer. BMC Genomics 2016;17:631

7. Zhu Y, Xu Y, Helseth DL, Jr., Gulukota K, Yang S, Pesce LL, et al. Zodiac: A Comprehensive Depiction of Genetic Interactions in Cancer by Integrating TCGA Data. J Natl Cancer Inst 2015;107

8. Niknafs YS, Pandian B, Gajjar T, Gaudette Z, Wheelock K, Maz MP, et al. MiPanda: A Resource for Analyzing and Visualizing Next-Generation Sequencing Transcriptomics Data. Neoplasia 2018;20:1144–9

9. Nicol JW, Helt GA, Blanchard SG, Jr., Raja A, Loraine AE. The Integrated Genome Browser: free software for distribution and exploration of genome-scale datasets. Bioinformatics 2009;25:2730–1

10. Freese NH, Norris DC, Loraine AE. Integrated genome browser: visual analytics platform for genomics. Bioinformatics 2016;32:2089–95

11. Chandrashekar D, Bashel B, Balasubramanya S, Creighton C, Ponce-Rodriguez I, Chakravarthi B, et al. UALCAN: A Portal for Facilitating Tumor Subgroup Gene Expression and Survival Analyses. Neoplasia 2017;19:649–58

12. Chandrashekar D, Karthikeyan S, Korla P, Patel H, Shovon A, Athar M, et al. UALCAN: An update to the integrated cancer data analysis platform. Neoplasia 2022;25:18–27

13. Tang Z, Li C, Zhang K, Yang M, Hu X. GE-mini: a mobile APP for large-scale gene expression visualization. Bioinformatics 2017;33:941–3

14. Zhu Y, Qiu P, Ji Y. TCGA-assembler: open-source software for retrieving and processing TCGA data. Nat Methods 2014;11:599–600

